# Evaluation of freshness and physicochemical characteristics of blankets of pirarucu (*Arapaima gigas*) bred in captivity at different ages in relation to the seasons

**DOI:** 10.1101/2021.08.04.455103

**Authors:** Krishna R. de Rosa, Alessandra A. da Silva, Wander M. de Barros, Anaqueli L. Pedroso, Maria Fernanda E. Ferreira, Luciana K. Savay-da-Silva, Lucia Aparecida de F. Mateus, Ernesto H. Kubota

**Author notes:** Corresponding author, (KRR). These authors contributed equally to this work (Alessandra A. da Silva, Anaqueli L. Pedroso, Maria Fernanda E. Ferreira).

## Abstract

This study aimed to evaluate the freshness and physicochemical characterization of pirarucu blankets at different ages bred in captivity in excavated tanks in northern Mato Grosso State, Brazil, in different seasons (rain and dry seasons). Four harvests were performed in two different tanks and five specimens obtained in each evaluated period, with animals aged 18 and 24 months, respectively, at the beginning of the experiment. The animals were slaughtered in a local establishment that had an inspection service following humane standards, and the blankets were later sent to analysis. The samples were analyzed for freshness via total volatile nitrogenous bases and physicochemical characteristics (pH in 24 h, water activity, water retention capacity, dripping loss, cooking loss, and shear force). There was a statistical difference for total nitrogenous volatile bases and shear force with higher values in the rainy season (24 month-of-old animals). Additionally, pH and water activity differed statistically at 24 h in the same period, albeit for 18-month-of-old animals. It was possible to conclude that climate variation affected the freshness and physicochemical characteristics of the pirarucu blankets while age did not.

## Introduction

Fish have a rich nutritional composition composed of highly digestible proteins and are a known source of polyunsaturated fats (e.g., omega-3) that help the cardiovascular system, fat-soluble vitamins A and D, and minerals such as phosphorus, iron, calcium, copper, selenium, and even iodine in salt-water specimens. Additionally, fish are widely known for having essential amino acids, including lysine, methionine, and cysteine [1, 2, 3, 4].

Notably, fish is a highly perishable food with high water activity, moisture content, and a source of nutrients. After slaughtering, numerous biochemical changes occur due to the speed of enzymatic reactions, triggering the loss of freshness [5, 6]. According to Jesus et al. [7], one factor associated with accelerated deterioration is cross-contamination due to internal contamination. Given this scenario and to provide a high-quality and safe product, it is pivotal for raw fish materials to be evaluated for freshness via total volatile nitrogen bases (TVNB), biogenic amines, and pH. Just as importantly, the physicochemical characteristics must also be determined, including water activity, water retention capacity, oxidative stability, and their influence on deterioration rates in fish are used to measure the intensity of the putrefaction process to ensure safe consumption [8, 9].

The pirarucu (*Arapaima gigas*) is the largest scaled fish in northern Mato Grosso State (Brazil) and inhabits the rivers of the Amazon basin as it prefers slow waters. The meat of this species is unique (boneless, with mild coloration and flavor, soft, and lean), and due to its condition of lung breathing, it must be slaughtered by concussion or hypothermia soon after capture [10, 11, 12].

Several factors, directly and indirectly, influence the productivity of a fish farm, with production management, fish morphometry, and region climatology being the most important. Based on the semi-intensive production system using excavated tanks whose water flow is usually continuous, properly managing the quality and quantity of this water source is essential in maintaining an environment conducive to aquaculture. Therefore, the seasons of the year affect fish farming, and rainfall alters the dynamics of shallow areas by modifying organic and inorganic material concentrations, heat dissipation, fish feeding habits, water turbidity and transparency, among others [13, 14].

Given the lack of research evaluating the influence of the seasons and the quality of this species, this study aimed to assess the freshness and physicochemical characteristics of the blanket of pirarucu of different ages and bred in captivity in excavated tanks during the rainy and dry seasons of the Amazon basin.

## Material and Methods

### Obtaining the raw materials

This study is part of a doctoral experiment and was carried out in strict accordance with recommendations of the Federal University of Santa Maria (UFSM - CAAE no. 09071018.4.0000.5346, CEP 3.210.927) and in partnership with the Longo fish farm whose procedures for breeding and slaughtering were duly approved by the Municipal Inspection Service (SIM 001/MT) and Unified Agricultural Health Care System (SUASA) for the sale of its products throughout the state of Mato Grosso, Brazil. All raw materials were obtained from fish farms (Tank A: 10°17′57.76“S and 54°56′29.34“W, altitude 342 m; and Tank B: 10°14′17“S and 54°56′1“W, altitude 261 m) and a slaughterhouse (10°14′17 “S and 54°56′1“W, altitude 261 m) in Peixoto de Azevedo (Mato Grosso State, the Amazon region).

Four harvests were performed in two excavated tanks and five specimens/tank/fish were collected, being five animals per collection in the rainy season (C1) in November/2017 for tank A (T1) and January/2018 for tank B (T2) and five animals in the dry season (C2) in June/2018 in T1 and July/2018 in T2, totaling 20 animals. The fish were 18, 24, 25, and 30 months old and had a total weight ranging from 7.69 to 15.67 kg and were 1.006 to 1.258 m long.

All specimens were bred on the fish farms and cultured identically from spawning to slaughter. The animals were fed in the morning with a dry artificial feed specific for fattening carnivorous fish (850 g kg^−1^ live weight) that had a maximum moisture content of 100 g kg^−1^, minimum crude protein content of 400 g kg^−1^, minimum ether extract of 100 g kg^−1^, maximum crude fiber content of 45 g kg^−1^, and maximum mineral content of 130 g kg^−1^. For both cases, the fish were kept in excavated tanks with the same dimensions: 6525 m^2^ in size (45 m wide, 145 m long, and 2 m deep in the water intake area and 3.5 m deep in the deepest part) and 17980 m^3^ of water.

Fish were harvested using trawl nets and the help of tractors on both sides of the tank and immediately transported to the SIM slaughterhouse located on the fish farm. Slaughter proceeded as follows: initially, the fish are stunned by ice water and slaughtered by bleeding, and after biometric analysis, they were processed to obtain the blankets (scaling, evisceration, beheading, fin removal, and bilateral maintenance by removing the spine). Each fish had its blankets placed in transparent polyethylene plastic packages, which are proper for food, and vacuum-sealed in an industrial sealer (R Baião, Minas Gerais, Brazil). The packages were then packed in styrofoam boxes with ice (±4 °C) and sent to the Meat and Fish Laboratory (LabCarPesc) of the Food Science and Technology College, Federal University of Mato Grosso, Cuiabá campus, where they were stored in an overnight cold chamber at 4 °C.

### Meteorological data

The study region (Peixoto de Azevedo) was founded in 1970 by the federal government through a colonization and occupation program of the central territory. This region has a peculiar biome that makes it part of the Brazilian Legal Amazon area that, in 2003, became part of the Amazon Portal Territory along with 15 other municipalities, such as Guarantã do Norte [15].

The Brazilian Amazon region has a distinctive climate with year-round high temperatures that vary in the dry (winter) and rainy (summer) periods. Moreover, this region is crucial given its vast forest areas that participate directly in the regional and global climate, intervening as a source of heat, humidity, and regulating the rainfall regime of both hemispheres [16].

Throughout the study period, the meteorological data of Peixoto de Azevedo (Köppen classification: humid tropical or sub-humid climate; Am) were provided by the National Institute of Meteorology (INMET) by the Automatic Weather Station of Guarantã do Norte installed in the Municipal Nursery (09°57′S and 54°53′W, altitude 320 m) because of its proximity to the fish farm (T1: 47.4 km away and T2: 38.1 km away). Among the information measured, we analyzed the average temperature (instantaneous, maximum, and minimum), average humidity (instantaneous, maximum, and minimum), and accumulated rainfall.

### Physicochemical and freshness analysis

The TVNB was determined in triplicate using a nitrogen distiller (model 4250, Thoth, São Paulo, Brazil) according to the adapted method of Savay-da-Silva et al. [17] and Normative Instruction no. 20 [18]. The pH levels in 24 h were directly measured in triplicate with a digital potentiometer (mPA-210, MS Tecnopon Instrumentation, São Paulo, Brazil) calibrated as described by Pregnolatto and Pregnolatto [19].

Water retention capacity (WRC) was analyzed using Whatman qualitative filter paper no. 1 (125 mm diameter, 50 × 50 cm acrylic plate, and specific weight of 10 kg) according to Hamm [20]. Drip loss (DL) was determined by hanging the blanket (100 g) in a net inside inflated plastic bags for 48 h at 4 °C in a cold chamber [21]. Cooking loss (CL) was performed in a water bath (N1040, Centaur, São Paulo, Brazil) at 85 °C for 1 h [22]. The water activity (Aw) was determined by dew-point measurements using a digital hygrometer (4TE, Aqualab, São Paulo, Brazil) [23], and shear force (SF) was measured with a digital texturometer (TA-XT Plus, Stable Micro Systems, Vienna Court, UK) programmed as follows [24]: return distance of 50 mm; return speed of 10 m/s; contact force of 5 g; pre-test speed of 1.0 mm/s; test speed of 2.0 mm/s; post-test speed of 10.0 mm/s; distance of 20.0 mm; and trigger force of 5 kg. All determinations were performed in triplicate except for texture analysis (quintuplicate for each replicate).

### Statistical analysis

For the physicochemical quality indices, data sets obtained from pretreatment screening were evaluated using one-way analysis of variance (ANOVA) and Tukey’s post hoc analysis (*p*>0.05 significance). Trends were only considered significant where means were different. Data normality was previously verified using the Shapiro-Wilk test, and all statistical analyses were performed by Statistica software (version 7.1) [25].

The effects of age and climate on the freshness and physicochemical characteristics of the blankets were evaluated using multivariate analysis techniques based on the similarity between objects (blankets) and attributes (variables). The variables were chi-square transformed to generate the similarity matrix [26], which was necessary because the variation scale of the measurements differed, and those with higher variation could determine the pattern if no transformation was done a priori. The Euclidean distance was then used as the similarity index.

PERMANOVA [27] was applied to evaluate the effects of age and climate (rainy and dry periods). This analysis answers whether the similarity matrix based on the measured variables presents a pattern and is associated with the previously established factors (age and period). The pattern was plotted by principal coordinate analysis (PCoA) in two dimensions (axes), and the correlation between PcoA axes and variables was estimated to assess which variables contributed the most to axis formation. Absolute values above 0.60 were considered significant. All analyses were performed in the R environment using the vegan package [28, 29]. The significance level adopted was 0.05.

## Results and discussion

### Meteorological data

The climatic data provided by the Guarantã do Norte automatic weather station is listed in Table 1. Temperatures remained high during the entire study period and showed a significant variation in humidity and accumulated rainfall.

**Table 1:**
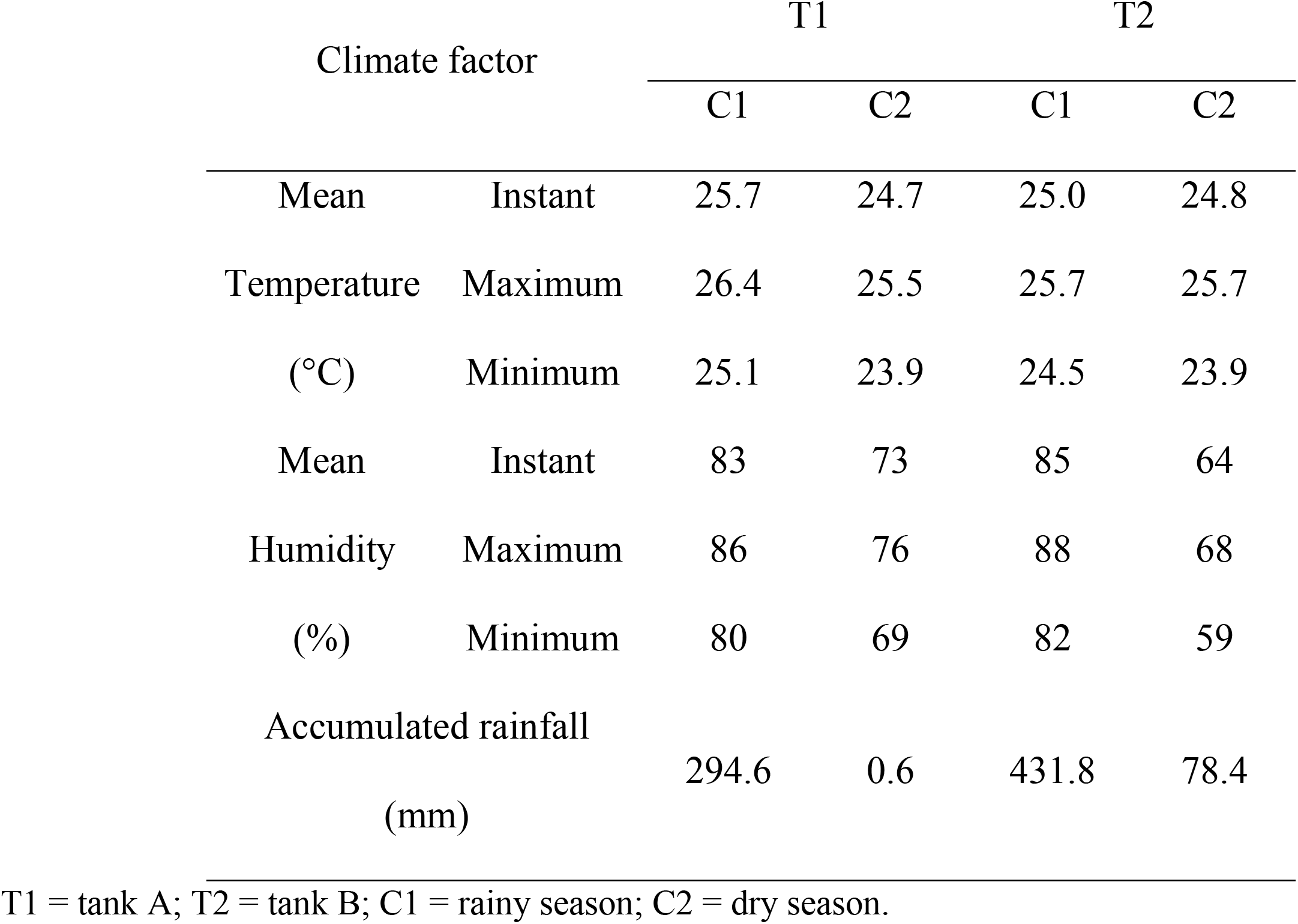
Climatic data from the Guarantã do Norte automatic weather station during the experimental period.

Santos [30] reported that the rainy period in Mato Grosso State lasts from November to January (minimum of 500-600 mm of rainfall), and the dry period occurs from June to August (maximum of 20-80 mm of rainfall). Hence, the data in Table 1 confirm that the accumulated rainfall reflects the period in question, albeit we removed the days in July 2018 (T2, C2) in which a subtropical jet stream brought a cold front from Patagonia and caused unseasonable rain.

Fish are actively affected by climate change because they are heterothermic, varying their metabolism and physiology to adapt to environmental temperature variations [31, 32]. Therefore, the species in the Amazon basin suffer from variations in the air-water relationship (solar radiation), soil leaching and siltation (flooding), changes in the feeding and nursery areas, low dissolved oxygen levels, and high temperatures [33]. According to Table 1, the average temperatures in the rainy season were higher than in the dry season, with maximum temperatures of 26.4 and 25.7 °C and a minimum of 24.5-25.1 and 23.9 °C, respectively. These findings corroborate the ideal temperature range described by Cyrino et al. [34] for tropical species aiming at thermal comfort for growth and reproduction (25 and 28 °C).

Moreover, research has proven there is a positive correlation between air and water temperatures, while rainfall and water temperatures are inversely correlated [35, 36, 37, 38]. Nonetheless, Matsuzaki et al. [39] and Millan [40] reported that water temperatures were higher during the rainy season (summer) in southern Brazil (São Paulo region), showing that the location affects the climatic condition and consequently the chemical composition of fish, which is also influenced by species, size, and sex of the animal [41, 42, 43, 44]. Therefore, water temperatures above the critical thermal maximum (e.g., 1 °C) can be lethal for fish, and this limit may vary from species to species, although it is usually higher in freshwater species. Alborali [45] studied rainbow trout in lakes and canals in Italy and reported that there was no change in fish behavior despite changes in water temperatures (from 11 to 18 °C), albeit oxygen and food consumption sharply increased.

### Analysis of freshness and physicochemical characteristics of pirarucu

There were no statistical differences for WRC, DL, and CL, while there were statistical differences and higher values for TVNB and SF (24-month-old fish) and pH in 24 h and Aw (18-month-old fish) in the rainy period (Table 2).

**Table 2.**
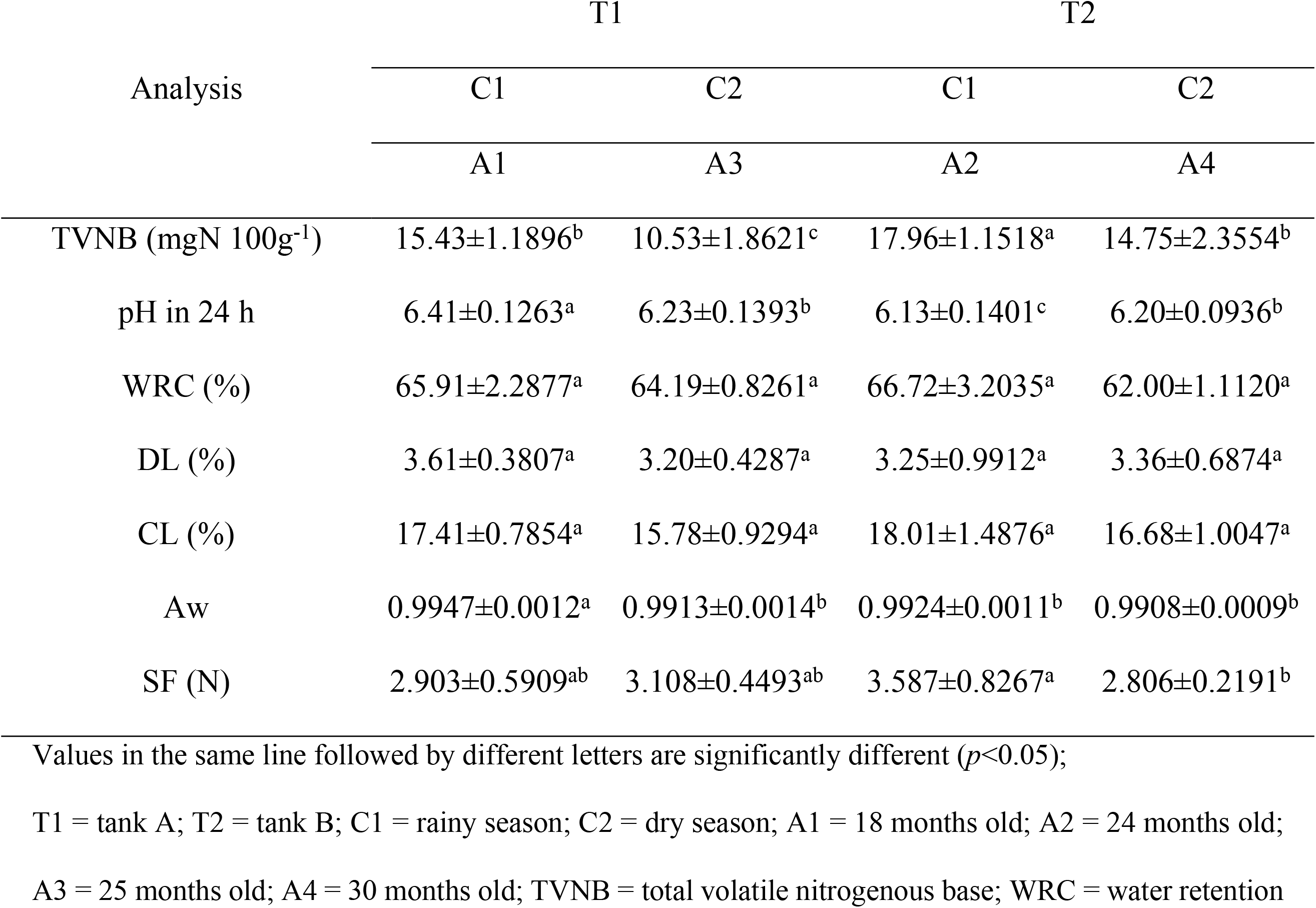
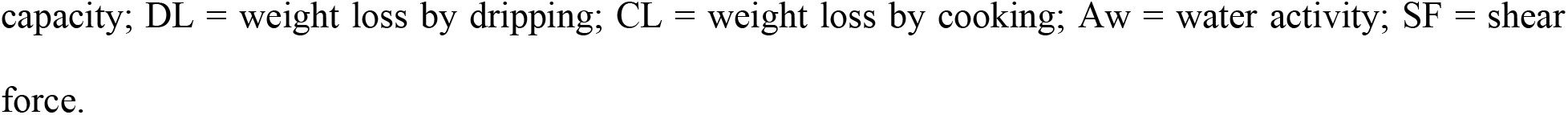
Physicochemical analysis of the blanket of pirarucu (*A. gigas*) bred in captivity. Results are expressed as mean ± standard deviation.

| Analysis | T1 |  | T2 |  |
| --- | --- | --- | --- | --- |
|  | C1 | C2 | C1 | C2 |
|  | A1 | A3 | A2 | A4 |
| TVNB (mgN 100g<sup>-1</sup>) | 15.43±1.1896<sup>b</sup> | 10.53±1.8621<sup>c</sup> | 17.96±1.1518<sup>a</sup> | 14.75±2.3554<sup>b</sup> |
| pH in 24 h | 6.41±0.1263<sup>a</sup> | 6.23±0.1393<sup>b</sup> | 6.13±0.1401<sup>c</sup> | 6.20±0.0936<sup>b</sup> |
| WRC (%) | 65.91±2.2877<sup>a</sup> | 64.19±0.8261<sup>a</sup> | 66.72±3.2035<sup>a</sup> | 62.00±1.1120<sup>a</sup> |
| DL (%) | 3.61±0.3807<sup>a</sup> | 3.20±0.4287<sup>a</sup> | 3.25±0.9912<sup>a</sup> | 3.36±0.6874<sup>a</sup> |
| CL (%) | 17.41±0.7854<sup>a</sup> | 15.78±0.9294<sup>a</sup> | 18.01±1.4876<sup>a</sup> | 16.68±1.0047<sup>a</sup> |
| Aw | 0.9947±0.0012<sup>a</sup> | 0.9913±0.0014<sup>b</sup> | 0.9924±0.0011<sup>b</sup> | 0.9908±0.0009<sup>b</sup> |
| SF (N) | 2.903±0.5909<sup>ab</sup> | 3.108±0.4493<sup>ab</sup> | 3.587±0.8267<sup>a</sup> | 2.806±0.2191<sup>b</sup> |

Values in the same line followed by different letters are significantly different ( $p < 0.05$ );
T1 = tank A; T2 = tank B; C1 = rainy season; C2 = dry season; A1 = 18 months old; A2 = 24 months old;
A3 = 25 months old; A4 = 30 months old; TVNB = total volatile nitrogenous base; WRC = water retention
capacity; DL = weight loss by dripping; CL = weight loss by cooking; Aw = water activity; SF = shear force.

The TVNB values (15.43, 10.53, 17.96, and 14.75) and pH in 24 h (6.41, 6.23, 6.13, and 6.20) were well below the limit established by Brazilian legislation (30 mgN 100g^−1^ and 7.0, respectively) [46]. The period in which the fish is harvested makes all the difference when analyzing the physicochemical composition and microbiological and nutritional quality. Moreover, the peculiarity of the species and individuality of each specimen [47, 48, 49] also play a critical role and interfere in the pH of the fish, causing it to increase due to the methods of capture, handling, and storage [50, 51].

As shown in Figure 1, there is an interaction between the age of the animals and climate period of the collections (F1;16= 2.21; *p*=0.104) for freshness and physicochemical composition, limiting the discussion of the effects of the factors separately (age: F1;16= 0.54; *p*=0.95; period: F1;16= 12.21; *p*=0.001). Furthermore, it was possible to observe that the freshness and physicochemical characteristics of the blankets were solely affected by the climate period (Figure 1).

**Fig 1.**
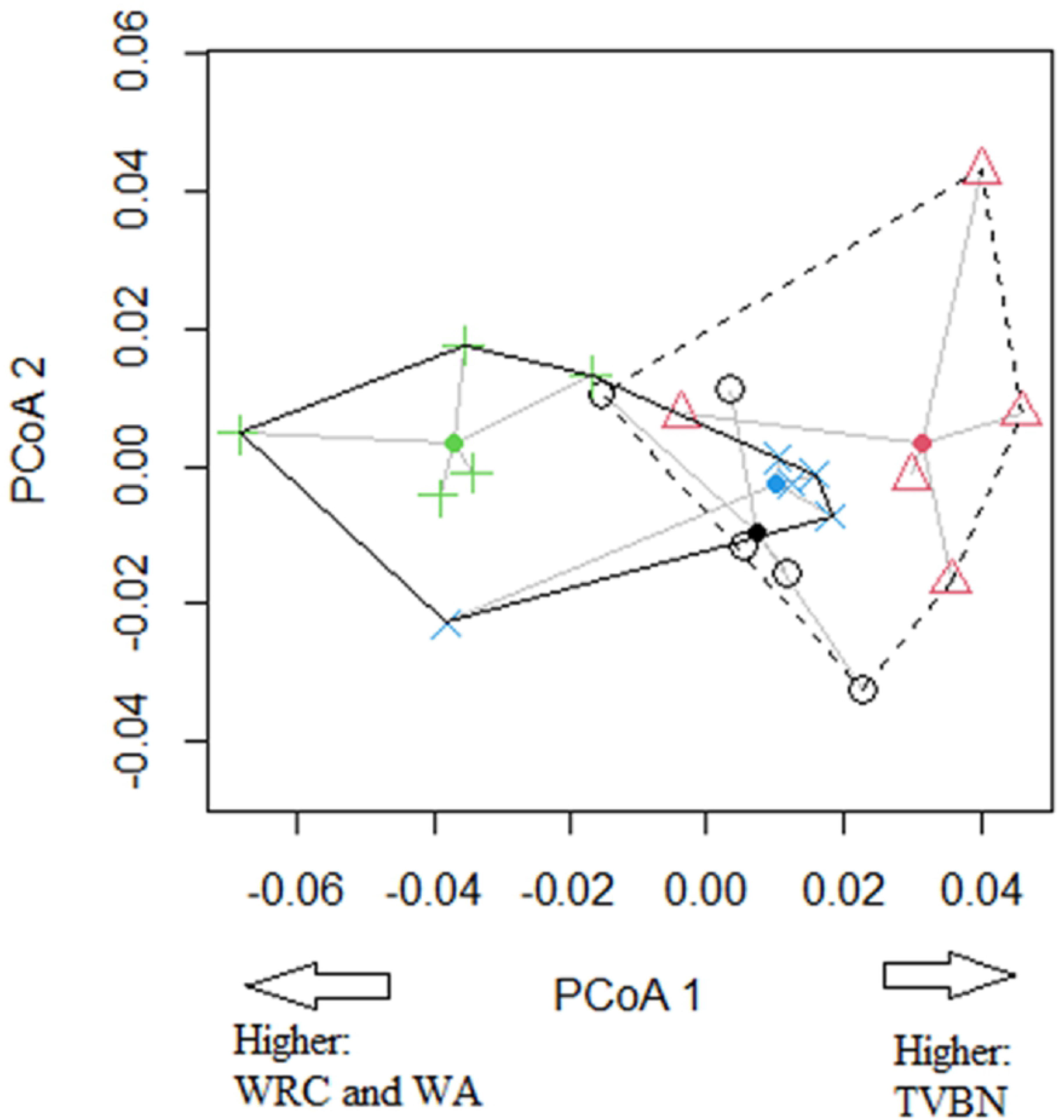
Effect of the age of the animals and the climatic period of the harvests on the freshness and physicochemical composition of the pirarucu blankets of the fish raised in captivity. (Circle = 18 months; triangle = 24 months; cross = 25 months; X = 30 months. Dotted polygon = rainy season; solid polygon = dry season). TVBN= total volatile base nitrogen; WRC= Water Retention Capacity; WA = Water Activity.

Axis 1 of the PCoA explained 52.93% of data variability, and this is associated with the interaction between the age and climate period and better separated the groups. It was possible to observe a positive correlation with axis 1 for some variables, implying that lower ages in rainfall have higher pH in 24 h, WRC, CL, and Aw values (variables that negatively correlated with axis 1) (Table 3).

**Table 3.** Correlations between axes 1 and 2 of the PCoA for freshness and physicochemical analysis of blanket of pirarucu bred in captivity.

| Analysis | Axes |  |
| --- | --- | --- |
|  | PCoA1 | PCoA2 |
| TVNB | 0.98585 | -0.00287 |
| pH in 24 h | -0.64847 | -0.22034 |
| WRC | -0.92852 | 0.228328 |
| DL | 0.189077 | -0.31319 |
| CL | -0.09234 | -0.7382 |
| Aw | -0.79009 | -0.16369 |
| SF | 0.042131 | 0.862495 |

TVNB = total volatile nitrogenous base; WRC = water retention capacity; DL = weight loss by dripping; CL = weight loss by cooking; Aw = water activity; SF = shear force.

Regarding tenderness, it is known that it is related to the composition of the muscle fibers, which are separated into three parts: red fibers (superficial part of the muscle in the subdermal layer), intermediate fibers, and white fibers (occupying over 90% of the muscle fibers and grant resistance to the fish) [52]. The concentration of each fiber varies according to the type of cut and fish species, and it can be indirectly quantified through the values obtained by the CIELab color system.

Data on the physicochemical profile of pirarucu are scarce and contradictory. Honorato et al. [53] evaluated 6.0 kg of pirarucu fillet steaks and reported pH values (6.27 ± 0.15) that were similar to the 25− and 30-months-old pirarucus analyzed herein for the dry season. However, Seering [54] characterized pirarucu slaughtered in the rainy period and found pH of 6.53 for pirarucu numbed in ice water and pH of 6.13 of fish that were not, also being similar to our findings for 18− and 24-month-old animals. Similarly, Jesus [55] reported TVNB values of 15.38 mgN 100g^−1^ for pirarucu belly, which is near our data for the rainy period and 18-month-old animals.

In the other parameters, most authors reported values above those described herein, although all of them are in accordance with the values required by Brazilian legislation for TVNB and pH levels exception for Jesus [55] in pirarucu belly (7.20). These high values, especially WRC and SF, may be associated with the origin of the raw material, which comes from older specimens and/or those that suffered stress during processing.

## Conclusions

The climatic variation influenced the freshness and physicochemical characteristics of the pirarucu blanket, while the age of the pirarucu did not. Higher total volatile nitrogenous bases, shear force, pH in 24 h, and water activity were obtained in the rainy period, albeit all parameters remained within limits mandated by Brazilian legislation, therefore proving that pirarucu blankets are a safe food.

## Acknowledgments

The authors would like to thank the Longo Fish farm for the partnership and for providing the raw materials. The authors would also like to thank Mariane Bittencourt Fagundes for greatly assisting in the statistical analysis of data and Atlas Assessoria Linguística for providing support with the English version of this manuscript.

## Supporting information

**Figure 1. The effects of age and climatic period of harvesting on the freshness and physicochemical composition of captive pirarucu blankets.** (Circle = 18 months; triangle = 24 months; cross = 25 months; X = 30 months. Dotted polygon = rainy season; continuous polygon = dry season. TVNB = total volatile nitrogenous bases; WRC = water retention capacity; Aw = water activity; CL = weight loss by cooking; DL = weight loss by dripping).

